# Field efficacy of guppies and pyriproxyfen (Sumilarv^®^ 2MR) combined with community engagement on dengue vectors in Cambodia: a randomized controlled trial

**DOI:** 10.1101/2020.05.15.097782

**Authors:** John Christian Hustedt, Dyna Doum, Vanney Keo, Sokha Ly, BunLeng Sam, Vibol Chan, Neal Alexander, John Bradley, Marco Liverani, Didot Budi Prasetyo, Agus Rachmat, Muhammad Shafique, Sergio Lopes, Leang Rithea, Jeffrey Hii

**Affiliations:** Epidemiology Department, Malaria Consortium, Phnom Penh, Cambodia; MRC Tropical Epidemiology Group, Department of Infectious Disease Epidemiology, London School of Hygiene and Tropical Medicine, London, United Kingdom; Department of Vector Control, Cambodian National Dengue Control Program, Phnom Penh, Cambodia; Department of Climate Change and Health, World Health Organization, Phnom Penh, Cambodia; Department of Entomology, US Naval Medical Research Unit-2, Phnom Penh, Cambodia

## Abstract

Evidence on the effectiveness of low-cost, sustainable biological vector control tools for *Aedes* mosquitoes is limited. Therefore, the purpose of this trial is to estimate the impact of guppy fish, in combination with the use of the larvicide Pyriproxyfen (Sumilarv^®^ 2MR), and Communication for Behavioral Impact (COMBI) activities to reduce entomological indices in Cambodia.

In this cluster randomized, controlled superiority trial, 30 clusters comprising of one or more villages each (with approximately 170 households) will be allocated, in a 1:1:1 ratio, to receive either a) three interventions (guppies, Sumilarv^®^ 2MR, and COMBI activities), b) two interventions (guppies and COMBI activities), or c) control (standard vector control).

Households were invited to participate, and entomology surveys among 40 randomly selected households per cluster were carried out quarterly. The primary outcome was the population density of adult female *Aedes* mosquitoes (i.e. number per house) trapped using adult resting collections. Secondary outcome measures include the House index, Container index, Breteau index, Pupae Per House, Pupae Per Person, mosquito infection rate, guppy fish coverage, Sumilarv^®^ 2MR coverage, and percentage of respondents with knowledge about *Aedes* mosquitoes causing dengue. In the primary analysis, adult female *Aedes* density and mosquito infection rates was aggregated over follow-up time points to give a single rate per cluster. This was analyzed by negative binomial regression, yielding density ratios.

The number of *Aedes* females was reduced by roughly half compared to the control in both the guppy and PPF arm (Density Ratio (DR)=0.54 [95% CI 0.34-0.85], p=0.0073), and guppy arm (DR=0.49 [95% CI 0.31-0.77], p=0.0021). The extremely low cost of including guppy rearing in community-based health structures along with the effectiveness demonstrated suggest guppies should be considered as a vector control tool as long as the benefits outweigh any potential environmental concerns. PPF was also highly accepted and preferred over current vector control tools used in Cambodia, however product costs and availability are still unknown.

**Author Summary:** Dengue is one of the most rapidly spreading mosquito-borne viral diseases in the world, is caused by bites of infected *Aedes* mosquitoes, and can sometimes lead to death. Cambodia has one of the highest per-capita incidence rates in Asia. Without a cure or routinely available efficacious vaccine, dengue control relies largely on reduction and avoidance of mosquitoes. In Cambodia, dengue mosquito control activities are focused on larviciding with temephos and pyrethroid based adulticide sprays to which *Aedes* have been shown to be increasingly resistant. This study was designed to evaluate novel biological vector control tools (guppy fish and a controlled release larvicidal matrix) utilizing an integrated vector management approach with community-based methods tailored to the local context. The results indicate that the tools resulted in a statistically significant reduction in immature and adult *Aedes* mosquito density. The interventions were accepted by and communities were willing to pay for them. The results suggest guppies are an ideal vector control tool as long as the benefits outweigh any potential environmental concerns. PPF was also highly accepted and preferred over current vector control tools used in Cambodia, however product costs and availability are still unknown.

## Introduction

Dengue is the most rapidly spreading mosquito-borne viral disease in the world, and is caused by bites of infected *Aedes* mosquitoes, principally *Aedes aegypti* [1]. Dengue is concentrated in the Asian region, which shoulders 70% of the global disease burden. Although a number of promising vaccine candidates are in preclinical and clinical development [2], innovative methods of genetic control of mosquitoes are being developed [3—6], however these interventions are unlikely to eliminate dengue on their own [7]. Therefore, traditional vector control will remain a key component of dengue control in the short and medium term.

In Cambodia, a total of 194,726 dengue cases were reported to the National Dengue Control Program (NDCP) between 1980 and 2008 [8]. However, the real number of cases and cost to society is estimated to be many times higher [9,10]. Previous work showed household water storage jars contained over 80% of *Ae. aegypti* larvae in Cambodia, and these jars became the main target for dengue vector control activities [11].

Since the early 1990s, NDCP has used the larvicide temephos (Abate^®^) to target large (200-400L) household water containers as the primary means of vector control [12]. This has continued despite tests published in 2001, 2007, and 2018 showing resistance of *Ae. aegypti* in several provinces across Cambodia [12–14]. Khun and Manderson (2007) concluded that “continued reliance on temephos creates financial and technical problems, while its inappropriate distribution raises the possibility of larvicide resistance.”[12] These problems led researchers to consider alternative control methods including chemical and biological substances (pyriproxyfen (PPF), and *Bacillus thuringiensis israelensis)* [1,12,15,16], jar covers [11], distribution of larvivorous copepods and fish [17–19]. The interventions that had the most effective results included the use of larvivorous fish and PPF[1,18].

The use of a larvivorous guppy fish *(Poecilia reticulata)* was evaluated in 14 Cambodian villages [17], and subsequently in a larger study of 28 Cambodian villages [18]. Results from the initial study conducted from 2006-2007 were encouraging as even with low coverage of guppies (in 56% of eligible containers one year after project commencement) there was a 79% reduction in *Aedes* infestation compared to the control area. Despite not having guppies, the smaller or discarded containers in the intervention area had 51% less infestation than those in the control area, suggesting a community-wide protective effect [17]. These results led the WHO and the Asian Development Bank (ADB) to fund a larger scale-up in 2010-2011 which included Communication for Behavioral Impact (COMBI) activities. At the end of the implementation period, an evaluation found that 88% of water jars, tanks, and drums contained guppy fish, suggesting successful establishment of breeding sites. In addition, the Container Index (the percentage of water holding containers infested with *Aedes* larvae or pupae) and the number of indoor resting adult females in the intervention area were near zero, while the control area had a Container Index of 30 [18]. Similarly encouraging results were found in Laos as a part of the same project, although many water containers in the implementation area were too small for guppy survival. This experience indicates that additional tools beyond larvivorous fish are required to target smaller water containers as well as hard-to-reach and cryptic breeding sites.

One potential solution to increase coverage of water containers in the communities is the use of PPF, a juvenile hormone analogue that interferes with the metamorphosis of juvenile *Aedes* mosquitoes, preventing their development. It can be used in small or contaminated containers unsuitable for larvivorous fish [20]. Studies of the efficacy of PPF in Cambodia showed inhibition of adult emergence (IE) greater than 87% for six months in 2003 [15], and IE above 90% for 20 weeks, and above 80% for 34 weeks in 2007 [1]. A slow-release PPF matrix release formulation (Sumilarv^®^ 2MR) has been developed and shown to be effective in Myanmar [21]. This new product only requires one distribution every six months (the entirety of the rainy season) so reduces operational costs as compared to temephos or *Bti* which have residual efficacy of 2-3 months [16,22].

Yet the efficacy of these measures, like other vector management approaches in the communities, is not only dependent on their entomological efficacy, but requires mobilization and coordination of resources to sustain behavior changes [23]. In particular, a key challenge for vector control in the communities is how local residents can be involved in and sustain vector breeding source reduction efforts [18]. Recent reviews indicate that a strong communication and behavior change approach, such as COMBI, has the potential to support vector management programs with very good outcomes [24,25]. For example, two new cluster randomized trials found that educational messages embedded in a community-based vector control approach were effective at reducing *Ae. aegypti* measured through entomological indices [26,27].

### Need for a trial

Although there is evidence suggesting the use of guppy fish can be beneficial in dengue vector control, recent reviews show there has never been a cluster randomized trial to evaluate their effect on mosquito indices [28]. This trial has the potential to inform the strategic application of community-based distribution of Pyriproxyfen and larvivorous fish in an outbreak, during inter-epidemic periods or for broad scale application. This trial will also be the first to our knowledge to evaluate the widescale use of the new Sumilarv^®^ 2MR product in the field. Furthermore, they have never been tested in combination. Our study is intended to fill these knowledge gaps.

### Hypothesis

This trial aims to demonstrate community effectiveness of guppies, PPF, and COMBI activities. The main hypotheses are:

1. Use of guppies, Sumilarv^®^ 2MR and COMBI activities will reduce numbers of *Aedes* mosquitoes, and their infection rates, more than guppies and COMBI alone, or standard vector control activities (such as larval control and information and education material dissemination during outbreaks) as assessed through entomology surveys;
2. COMBI activities will improve the community’s knowledge, attitudes, and behavior related to water use and vector borne disease prevention (such as burning or burying discarded containers, cleaning the environment around the house, and sleeping under a bednet) as assessed through baseline/endline surveys and Focus Group Discussions (FGDs);
3. Guppies and pyriproxyfen will be acceptable among the target villages as assessed by an endline survey and FGDs.

## Methods

This study followed the Consolidated Standards of Reporting Trials (CONSORT) guidelines [29] (S1 Table).

### Study design and setting

The study is designed as a cluster randomized, controlled trial with three arms. The study has 30 clusters in the province of Kampong Cham, where each cluster is a village or group of villages with on average 170 households (range 49-405) or 757 individuals (range 250-1769). The rainy season runs from April to November, and the peak dengue season is from May to July. The clusters were selected in areas which had *Aedes* infestation in the past. To minimize potential spillover effects, clusters had to be at least 200 meters from the nearest household outside the cluster since *Ae. aegypti* in this region have an average flight range of 50-100m [30]. Every house within the cluster boundaries was invited to participate in the trial.

### Interventions

Selected villages were randomized into one of three study arms (**Table 1**). Reasons for selecting the interventions for each arm are described above and in more detail in the study protocol [31]. The total trial period for the interventions was 11 months (S1 Figure).

**Table 1:**
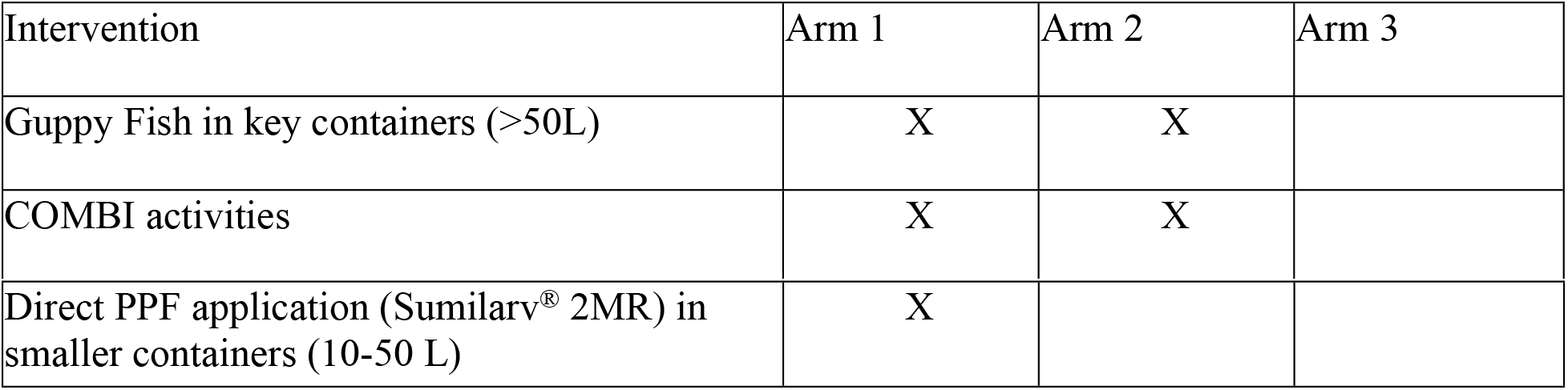
Interventions randomized to each study arm

#### Guppies

Two guppy fish (*Poecilia reticulata*) were placed into each water container greater than 50L in intervention villages (Arms 1 and 2). This is based on larval consumption of guppies determined by Seng et al. [17] and past experiences using guppies in vector control in Cambodia [18]. The guppies were sourced from the original NDCP colony, which was started from guppies found in a rural waterway near Phnom Penh roughly fifteen years earlier. The guppy fish were distributed after the baseline activities through a local community network managed by provincial government authorities [31]. CHWs were provided two jars for rearing. Each month CHWs conducted visual checks and ensured all their assigned households have guppies in all large containers and replaced them if necessary.

#### Pyriproxyfen Matrix Release (Sumilarv^®^ 2MR)

The product contains pyriproxyfen incorporated in an ethylen copolymer resin disk, and the PPF is gradually released from the polymer material until it reaches an equilibrium state of the dissolved active ingredient with that in the matrix formulation [32]. Each device is designed to provide coverage for 40 L of water, and can be cut into smaller sizes for smaller containers [31]. PPF devices were distributed to containers of size 10-50 liters at the beginning of the trial and replaced after 6 months. Additional devices were left at the HC for CHWs to distribute during their monthly monitoring visit if some were lost or needed to be replaced. The exceptional safety of PPF is reflected in WHO’s statements that it is “unlikely to present acute hazard in normal use”, “pyriproxyfen does not pose a carcinogenic risk to humans”, and PPF “is not genotoxic.” As a result of its efficacy, The WHO Pesticide Evaluation Scheme has recommended the use of pyriproxyfen for mosquito control [33]. Animal models suggest a very favorable mammalian toxicity profile, and extremely low risk for humans using this product [30].

#### Communication for Behavioral Impact Activities

A rapid formative assessment consisting of FGDs and In-Depth Interviews (IDIs) regarding knowledge, attitudes, and behaviors of community members was completed. The results formed the basis of well-informed COMBI interventions and were used in a message and material development workshop held with key community and district stakeholders [31]. Two days of training were given to CHWs on communication and facilitation skills, roles and responsibilities, and community participation following which they took the lead role in conducting health education sessions twice every month in their community [31]. Monthly meetings were also conducted with CHWs to assess progress, address issues and challenges, and provide them continuous training.

### Adherence

In order to improve adherence to the intervention protocols, CHWs performed monthly monitoring checks on each household within the intervention arms, and entomology surveys recorded the presence or absence of each intervention in containers [31]. Project staff also randomly visited CHWs and intervention households to confirm the reliability of data provided.

### Primary Outcome Measures

The primary outcome measure is the population density (i.e. number of mosquitoes per unit of time spent aspirating) of adult female *Aedes* trapped using adult resting collections.

### Secondary Outcome Measures

The secondary outcomes for the trial include:

1. Dengue virus infection rate in adult female *Aedes* mosquitoes
2. House index (HI): Proportion of houses surveyed positive for *Aedes* larvae and/or pupae in any water container
3. Container index (CI): Proportion of surveyed containers containing *Aedes* larvae and/or pupae
4. Breteau index (BI): Number of containers positive for *Aedes* larvae and/or pupae per 100 houses surveyed
5. Pupae Per House (PPH): Number of *Aedes* pupae per household
6. Pupae Per Person (PPP): Number of *Aedes* pupae per person
7. Guppy fish coverage: proportion of eligible water containers with ≥1 guppy fish
8. Sumilarv^®^ 2MR coverage: proportion of eligible water containers with ≥1 MR resin disc
9. Percentage of respondents with knowledge about *Aedes* mosquitoes causing dengue

### Sample Size

The guppy fish and pyriproxyfen interventions were assessed by four entomology surveys. A sample size of 10 clusters per arm and 40 HHs per cluster for the survey was devised using the Hemming and Marsh method [34] assuming a mean of 0.1 adult female resting *Aedes* per household in the intervention arms compared to 0.25 in the control arm for each collection based on previous studies. The households were randomly selected each collection. The intracluster correlation (ICC) was assumed to be 0.01 based on previous studies [31]. Additionally, a sensitivity analysis was conducted up to the median value of ICCs for outcome variables (0.03) as found by an analysis conducted by Campbell et al. [35]. Our analysis determined that ICC values between 0.01 to 0.03 would have 91 to 75% power, respectively.

The impact of COMBI activities in the communities was evaluated through Knowledge, Attitudes, and Practice (KAP) surveys. A sample size of 10 clusters per arm and 20 HHs per cluster was devised, again using the Hemming and Marsh method [34], assuming a 22.5% change in KAP indicators from 40% to 62.5% in intervention villages and no change in the control villages over the course of one year [31].

### Allocation

Clusters were randomly assigned with a 1:1:1 allocation through a public randomization process. Village chiefs from all clusters and HC chiefs from all HCs were invited to a central point along with local and national authorities, where allocation took place. Allocation concealment was accomplished by having each cluster representative choose one folded up paper with a printed label referring to arm one, two, or three.

### Data collection methods

Data were collected at 0, 4, 8, 12 months post-intervention, unless otherwise mentioned. The timing was also meant to capture data over different season (e.g. heavy rain, light rain, and dry seasons). The project employed the following methods:

#### Entomology

A baseline survey was conducted prior to start of interventions. An endline survey was conducted one year after the baseline. Two additional surveys during the dry season (4 months post intervention) and light rain (8 months post intervention - peak dengue season) were also conducted. The survey methodology was developed following the WHO guidelines for entomological collections [36] and detailed in the study protocol[31]. The survey team also completed a rapid assessment tool (Premise Condition Index) (PCI) [37] to identify whether the scores can predict household risk for *Ae. aegypti* infestation [38].

#### Knowledge, Attitudes, and Practices

KAP surveys were conducted at the same time as baseline and endline entomology surveys [31]. The secondary outcome measure included was whether participants knew dengue is transmitted by mosquitoes of the genus *Aedes,* rendered in Khmer as “kala”, meaning feline or tiger.

#### Community Health Worker monthly monitoring

The coverage of guppy fish and PPF Sumilarv^®^ 2MR were assessed by ocular inspection of water containers via entomology surveys and the CHW monthly reporting form as described in the adherence section. Coverage is expressed as percentage of containers with at least two guppy fish or one Sumilarv^®^ 2MR of the total households or containers examined.

#### Climate

General climate data (rainfall, temperature and humidity) were recorded at one of the intervention health centers using a rain gauge and a Hobo onset data logger. All villages have virtually the same climate.

#### Data Management

The first two entomology surveys and the first KAP survey were recorded on paper, and double data entry was performed using EpiData (EpiData Association, Denmark) by an experienced data processing company. Due to factors including budget, timeliness, and need for data cleaning, the subsequent two entomology surveys and final KAP survey were recorded electronically on Samsung tablets (Samsung Group, South Korea) and uploaded to ONA servers (ONA, USA).

### Mosquito flavivirus infection

Adults female *Aedes* mosquitoes were pooled together by cluster with a maximum of 10 per pool, and an expected minimum infection rate of 3-7% based on other studies [39,40]. Flavivirus detection in adult female mosquitoes followed the protocol set out by Pierre et al. [41] using a set of universal oligonucleotide primers. Samples identified as positive for flavivirus were then put into a rapid assay for detecting and typing dengue viruses [42]. All pools had positive and negative controls to ensure the tests were working properly.

#### Statistical Methods

All statistical analyses were performed in R version 3.5.0 (Murray Hill, New Jersey) and Stata^®^ version 14.2 (College Station, Texas).

#### Primary Outcome

Adult female *Aedes* density was summed over follow-up time points to give a single rate per cluster. This was analyzed by negative binomial regression using the number of adults as the response, and the logarithm of the sampling effort (that is, person-time spent aspirating) as an offset. Hence, this analysis yielded density ratios.

#### Secondary Outcomes

None of the mosquito pools tested were positive for dengue virus; consequently, the minimum infection rate was 0%. The most commonly used entomological indexes (BI and PPP) are reported here, where correlated indices (CI, HI, and PPH) are listed in the supplementary tables (Table S4.1).

### Data Monitoring

In accordance with the recommendations of Grant et al., we did not establish a Data Safety Monitoring Board for this study as it is not a “clinical trial evaluating a therapy with a mortality or irreversible morbidity endpoint” [43]. However, a Technical Steering Committee (TSC) was established which met at least every six months and addressed any concerns that arose [31]. Participants were told to report any adverse events directly to project staff or CHWs and seek medical attention immediately. CHW monthly monitoring forms include a line to report any adverse events that have taken place. Any report of harm or adverse events was reported directly to the TSC.

### Access to Data

All co-principal investigators and partners were given access to the cleaned data sets without identifiers, which were stored on the Malaria Consortium Sharepoint site and were password protected. The final anonymized dataset will be stored in the Cambodian National Centre for Parasitology, Entomology, and Malaria Control central repository and the final cluster level dataset used for the analysis in the results section is attached as supporting material (S1 Dataset). Entomological specimens are stored for two years at Malaria Consortium offices should other researchers be interested in accessing them.

### Ethical Approval and Consent to participate

Ethical clearance for this trial was received by the Cambodian National Ethics Committee for Health Research on Oct 9th, 2014 (ethics reference number 0285). Additionally, ethics approval was received from the London School of Hygiene and Tropical Medicine Observational / Interventions Research Ethics Committee (ethics reference number 8812). CHWs explained the trial and received informed consent from the head of the household before providing the interventions [31]. Those who were illiterate or otherwise could not sign their name were given the option of giving their thumb print. All village and respondent names were deleted to ensure no identifying information was included. Data from surveys were stored in a password-protected computer. All qualitative data were collected in concordance with the guidelines of the Code of Ethics of the American Anthropological Association (AAA) [44].Results

### Baseline Results

In the baseline results the control arm had a slightly larger number of houses/people than in intervention arms (Table 2). The sex and age distribution of household heads was similar between the three arms. The mean number of containers, positive containers, BI, and PPP at cluster level were all larger in the guppy only arm (arm 2) than others, while the mean number of adult *Aedes* females per cluster was similar between arms.

**Table 2:** Baseline summary measures of containers, houses, and people per cluster

|  | Control | Guppies | PPF + Guppies |
| --- | --- | --- | --- |
| Number of Clusters | 10 | 10 | 10 |
| Number of houses | 2016 | 1641 | 1435 |
| Number of people | 8475 | 7542 | 6700 |
| Number of houses surveyed | 400 | 400 | 400 |
| Percentage of Male Household Heads (Range) | 22 (10-45) | 23 (10-32) | 20 (10-35) |
| Median Age of Household Head (Range) | 42 (17-78) | 42 (18-84) | 45 (18-88) |
| Mean Number of Containers Per Cluster (Range) | 154 (121-190) | 186 (160-219) | 165 (110-213) |
| Mean Number of Positive Containers Per Cluster* (Range) | 24.7 (18-62) | 36.5 (18-62) | 27.7 (11-69) |
| Mean Breteau Index Per Cluster (Range) | 62 (20-115) | 91 (45-155) | 69 (28-173) |
| Mean Pupae Per Person (Range) | 0.9 (0.2-2.7) | 4.0 (0.2-17.1) | 1.1 (0.5-2.3) |
| Mean Adult *Aedes* Female Density Per Cluster (Range) | 10 (1-15) | 9 (3-24) | 11 (2-20) |

\*Positive is defined as having either *Aedes* pupae or larvae in the container.

### Primary Outcome

Over the intervention period, the population density of adult female *Aedes* was significantly less in both the guppy + PPF arm (Arm 1) (Density Ratio (DR)=0.54 [95% CI 0.34-0.85], p=0.0073), and guppy arm (Arm 2) (DR=0.49 [95% CI 0.31-0.77], p=0.0021) relative to control (Arm 3). However, the difference between the two intervention arms was not significant (DR=1.10 [95% CI 0.69-1.74], p=0.6901) (Table 3). The mean number of adult *Aedes* females was the highest in the light rain season and lowest in the rainy season. (Fig 1).

**Fig 1:**
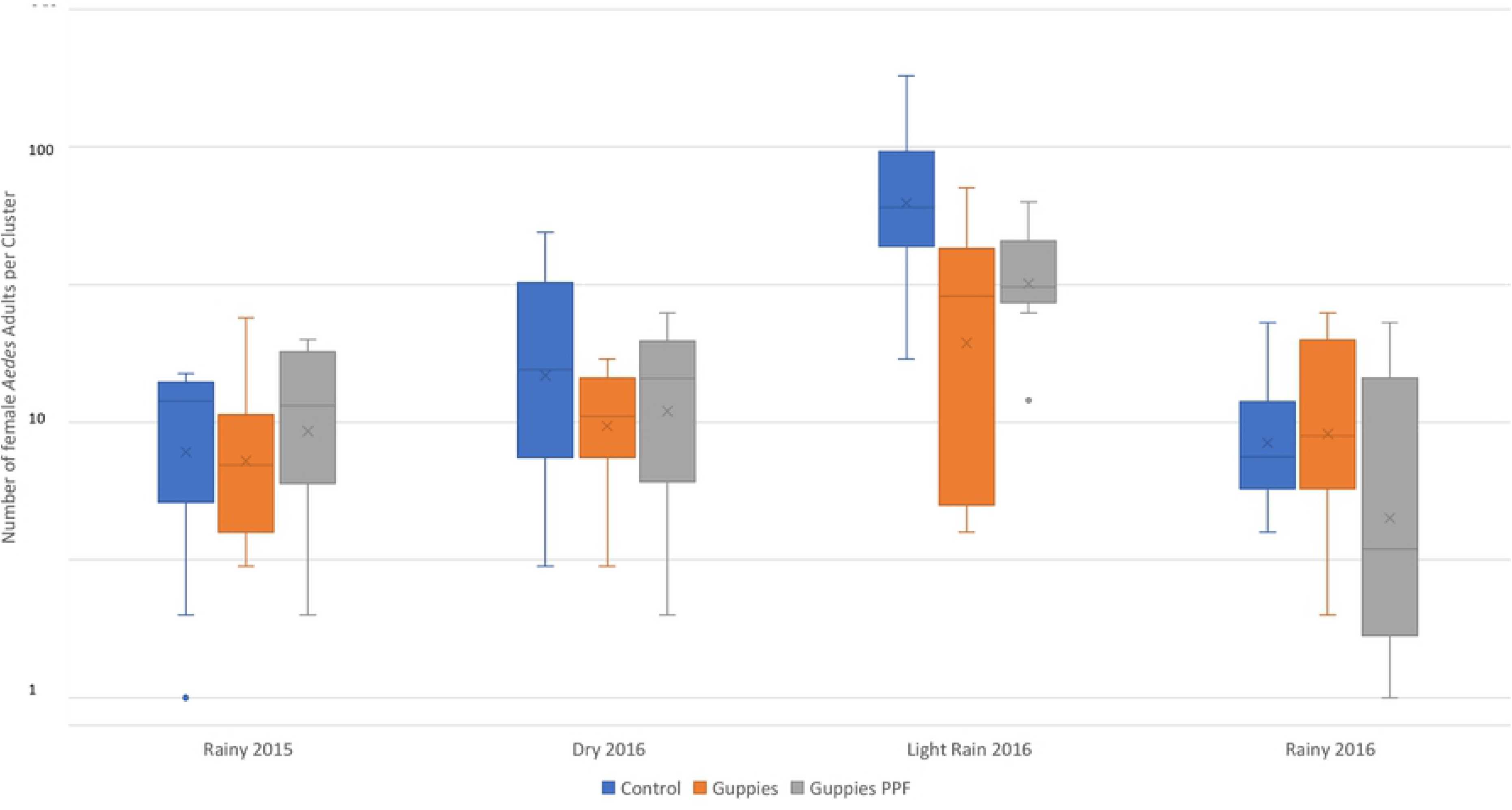
Box plots showing mean number of adult Aedes females per household by arm and season, October 2015 – October 2016

**Table 3:** Mean population density of adult female *Aedes* trapped using adult resting collections per cluster by arm and survey

|  | Control | Guppies | Guppies + PPF |
| --- | --- | --- | --- |
| Baseline (Range) | 10 (1-15) | 9 (3-24) | 11 (2-20) |
| Dry Season (Range) | 20 (3-49) | 11 (3-17) | 14 (2-25) |
| Light Rain (Range) | 75 (17-181) | 29 (4-71) | 35 (12-63) |
| Heavy Rain (Range) | 10 (4-23) | 12 (2-25) | 8 (1-23) |
| Total (Range) | 35 (3-181) | 17 (2-71) | 19 (1-63) |
| Density Ratio (95% CI), p-value* | 1 (Ref) | 0.49 (0.31-0.77),<br>p=0.0021 | 0.54 (0.34-0.85),<br>p=0.0073 |
| Density Ratio (95% CI), p-value* | ** | 1 (Ref) | 1.10 (0.69-1.74),<br>p=0.6901 |

\*The ratios do not include the baseline data
\*\*The ratio is not given here as it would be redundant
The trapping time was 10 minutes per house

### Secondary Outcomes

No adult female *Aedes* mosquitoes in any arm were found to be positive by PCR for dengue virus (n=280 pools). The most commonly used entomological indexes (BI and PPP) are reported here, where correlated indices (CI, HI, and PPH) are listed in the supplementary tables (S2 Table).

#### Breteau Index

Over the intervention period, the BI was significantly less in both the guppy + PPF arm (Arm 1) (DR=0.64 [95% CI 0.50-0.85], p=0.0016), and guppy arm (Arm 2) (DR=0.63 [95% CI 0.48-0.82], p=0.0006) relative to control (Arm 3). The difference between the two intervention arms was not significant (DR=0.97 [95% CI 0.73-1.27], p=0.7982) (Table 4). The biggest difference between arms was seen during the dry and light rain or rainy seasons (Fig 2).

**Fig 2:**
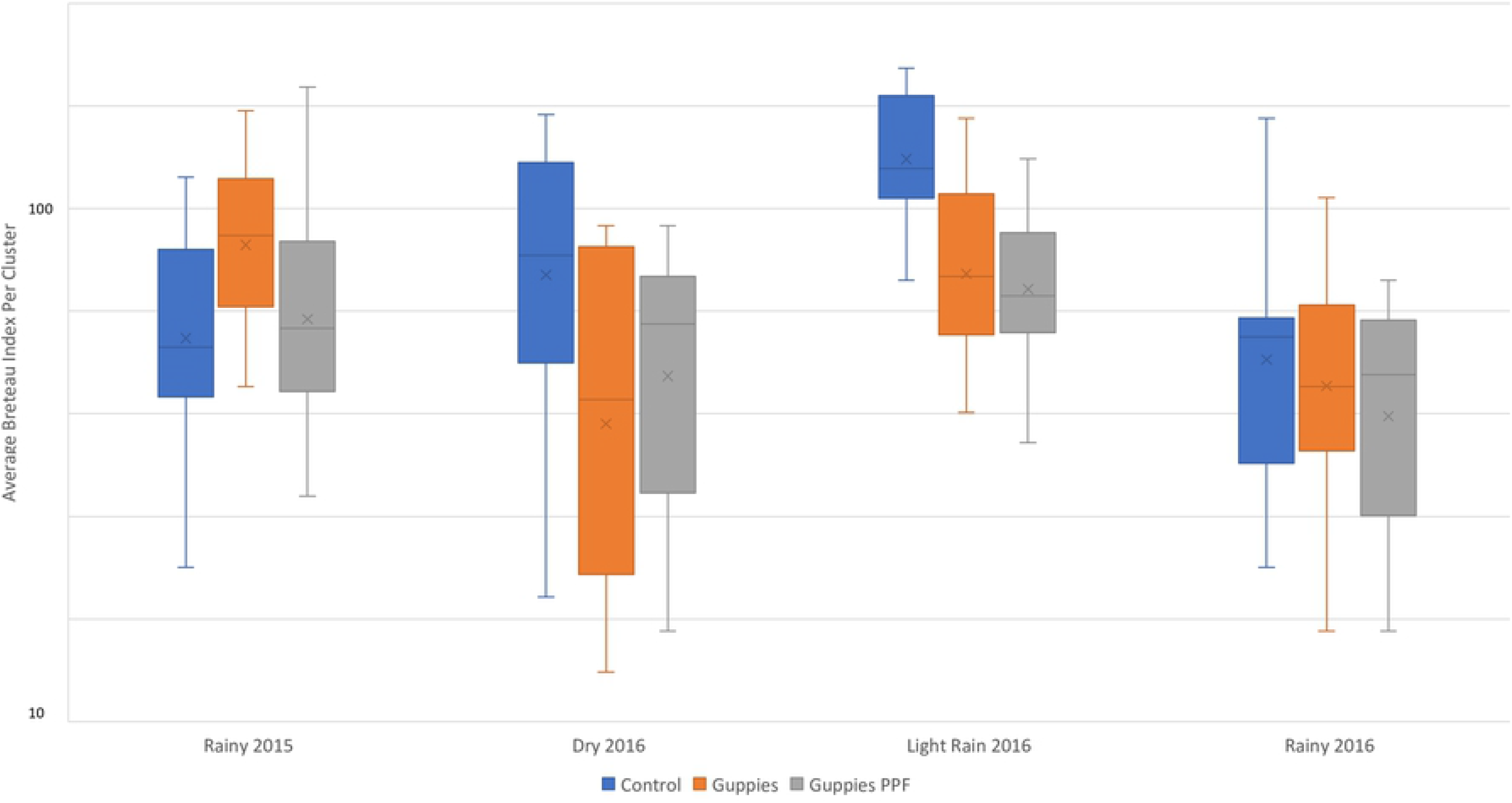
Box plots showing Breteau index by arm and season, October 2015 – October 2016

**Table 4:** Immature *Aedes* indices per cluster by arm and survey

**Fig 2: Box plots showing Breteau index by arm and season, October 2015 – October 2016**
| **Table 4: Immature *Aedes* indices per cluster by arm and survey** |  |  |  |
| --- | --- | --- | --- |
|  | **Breteau Index** |  |  |
|  | Control | Guppies | Guppies + PPF |
| Baseline (Range) | 62 (20-115) | 91 (45-155) | 69 (28-173) |
| Dry Season (Range) | 88 (18-153) | 48 (13-93) | 54 (15-93) |
| Light Rain (Range) | 130 (73-188) | 81 (40-150) | 74 (35-125) |
| Heavy Rain (Range) | 58 (20-150) | 51 (15-105) | 45 (15-73) |
| Total (Range) | 92 (18-188) | 60 (13-150) | 58 (15-125) |
| Density Ratio (95% CI), p-value* | 1 (ref) | 0.65 (0.50-0.85), p=0.0016 | 0.63 (0.48-0.82), p=0.0006 |
| Density Ratio (95% CI), p-value* | ** | 1 (ref) | 0.97 (0.73-1.27), p=0.7982 |
|  | **Pupae Per Person** |  |  |
|  | Control | Guppies | Guppies + PPF |
| Baseline (Range) | 0.9 (0.2-2.7) | 4.0 (0.2-17.1) | 1.1 (0.5-2.3) |
| Dry Season (Range) | 1.0 (0.1-3.3) | 0.3 (0-0.9) | 0.7 (0-1.7) |
| Light Rain (Range) | 2.2 (0.5-7.0) | 1.2 (0.1-3.3) | 0.60 (0-1.4) |
| Heavy Rain (Range) | 0.7 (0.1-2.1) | 0.6 (0.1-2.9) | 0.7 (0-1.8) |
| Total (Range) | 1.3 (0-7.0) | 0.7 (0-3.3) | 0.7 (0-1.8) |
| Density Ratio (95% CI), p-value* | 1 (ref) | 0.56 (0.35-0.91), p=0.0193 | 0.52 (0.32-0.84), p=0.0075 |
| Density Ratio (95% CI), p-value* | ** | 1 (ref) | 0.92 (0.60-1.49), p=0.7385 |

\*The ratios do not include the baseline data
\*\*The ratio is not given here as it would be redundant

#### Pupae Per Person

Baseline results show significantly higher PPP in the guppy arm (Arm 2) than the other arms (Fig 3). Over the intervention period, the PPP was significantly less in both the guppy + PPF arm (Arm 1) (DR=0.56 [95% CI 0.35-0.91], p=0.0193), and guppy arm (Arm 2) (DR=0.52 [95% CI 0.32-0.84], p=0.0075) relative to control (Arm 3). The difference between the two intervention arms was not significant (DR=0.92 [95% CI 0.60-1.49], p=0.7385) (Table 4).

**Fig 3:**
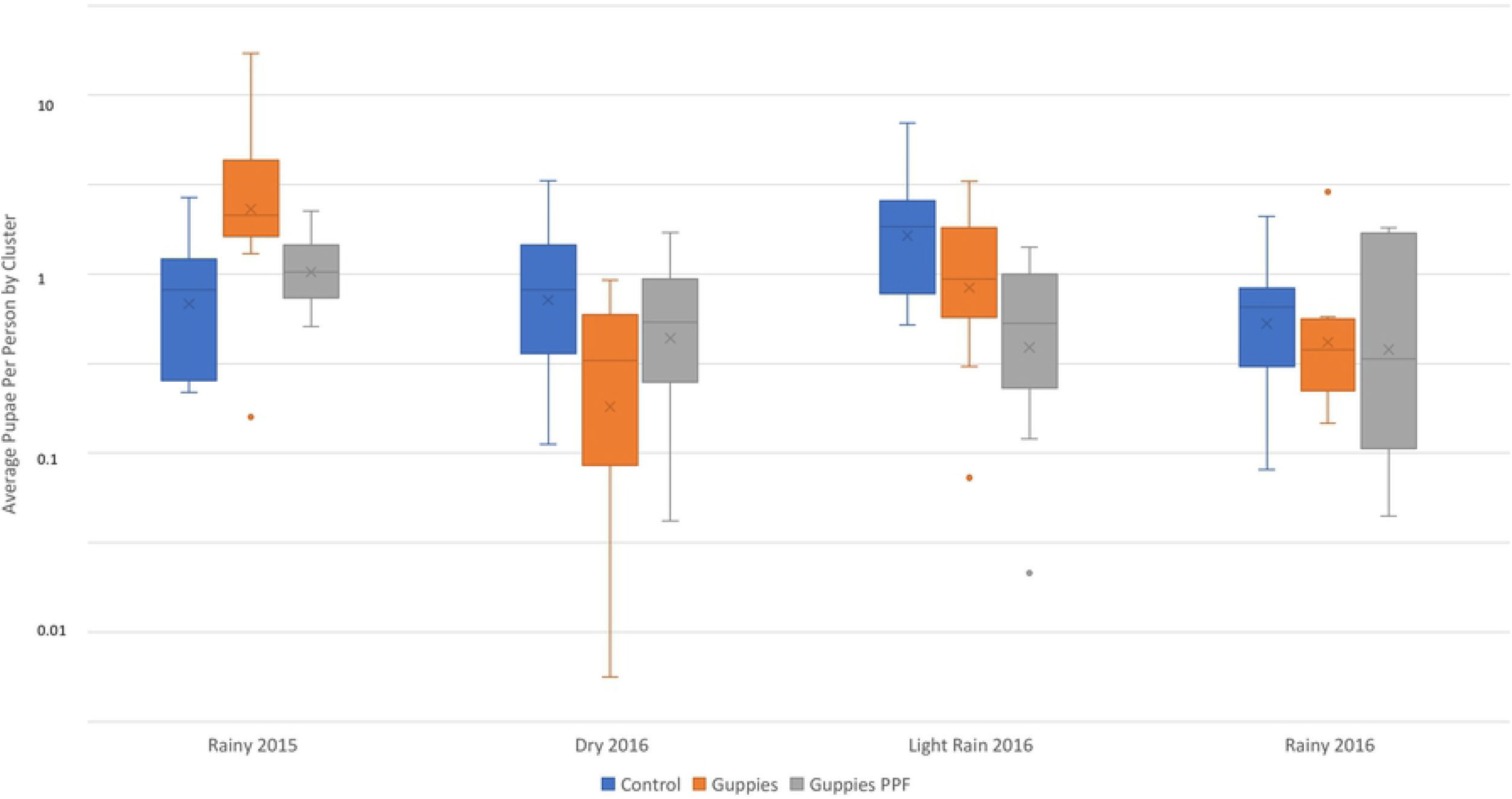
Box plots showing pupae per person by arm and season, October 2015 – October 2016

#### Knowledge, Attitudes, and Practice Survey

The secondary outcome related to the KAP survey is reported here, while the full data set from the KAP survey is in the supplementary files (S2 Appendix). High levels of knowledge that dengue is transmitted by *Aedes* mosquitoes were reported at baseline among all arms (range 95.5-98%). Endline surveys showed 100% of participants with this knowledge. Ratios of increased knowledge between baseline and endline were not significantly different between arms with the guppy + PPF arm (Arm 1) (RR=0.99 [95% CI 0.86-0.1.14], p=0.915), and guppy arm (Arm 2) (RR=1.01 [95% CI 0.87-1.16], p=0.943) relative to control (Arm 3) (S2 Table).

#### Coverage of Guppy Fish and Sumilarv^^®^^ 2MR

Coverage of guppy fish (proportion of eligible water containers with ≥1 guppy fish) before replacement in Arm 2 rose to nearly 80% after one month and stayed close to 70% for most of the intervention period (Fig 4). However, in Arm 1 PPF coverage (proportion of eligible water containers with ≥1 Sumilarv*^®^* MR) rose to 80% after two months and stayed high until dropping in March, after which continued health education messages increased coverage back to near 70-80%. Guppy coverage in Arm 1 was notably lower (near 50%) until guppy use was emphasized in March, after which it increased dramatically and then dropped off back to around 50%.

**Fig 4:**
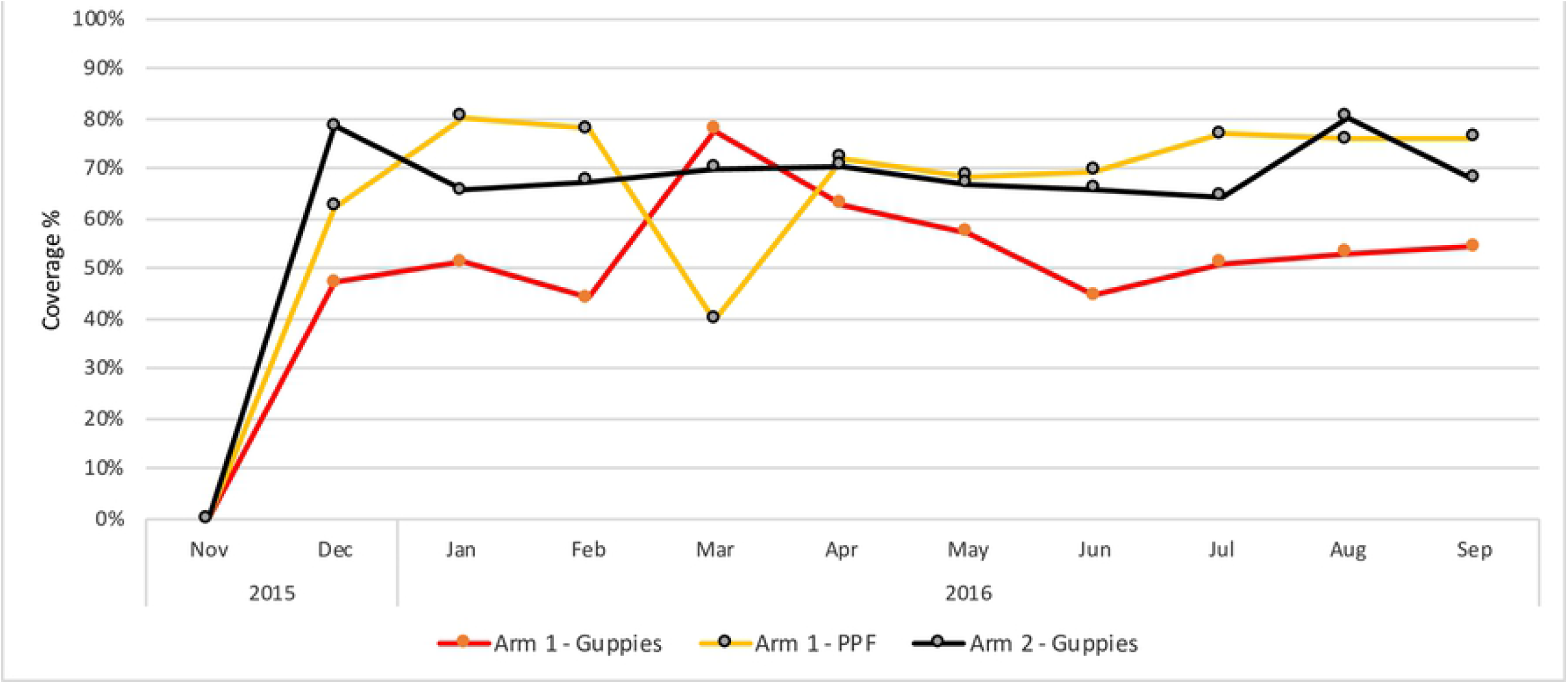
Coverage of guppies and PPF in intervention villages by month, November 2015 - September 2016

#### Climate

The average maximum daily temperature in the shade decreased from 34.4°C in the dry season to 31.3° C in the light rain season. The average relative daily humidity and monthly rainfall increased from 60.0% and 10.7 millimeters to 75.2% and 139 millimeters from the dry to light rain season, respectively (Fig 5). The rainy season saw much larger amounts of rainfall (near 300 millimeters per month) than all other seasons.

**Fig 5:**
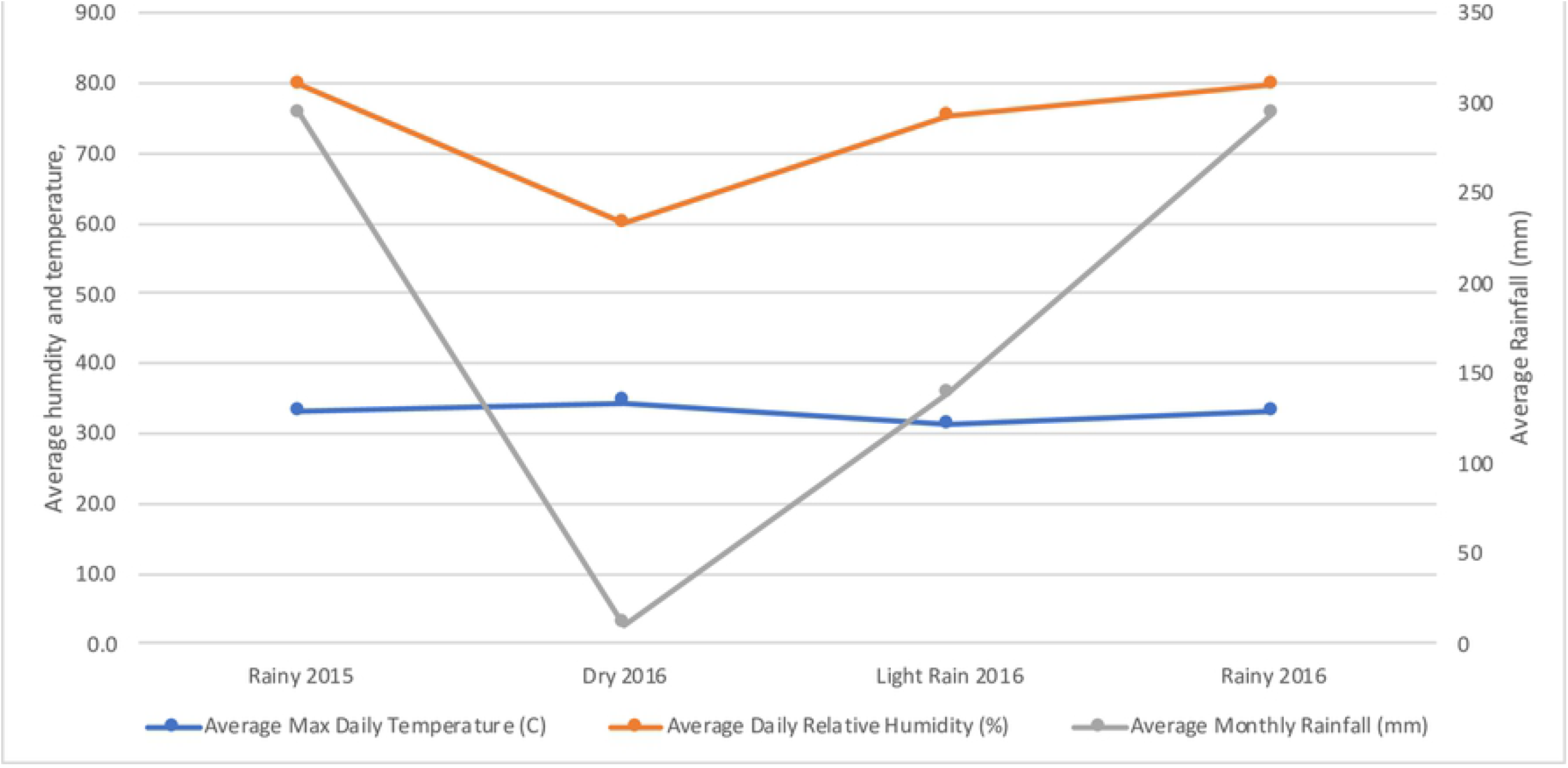
Average maximum daily temperature, relative humidity, and rainfall

### Adverse Events

No adverse events, harms, or unintended effects were recorded during the trial.

## Discussion

Guppies, whether or not in combination with PPF, were able to decrease the number of *Aedes* females (DR=0.49-0.54) and PPP (DR=0.52-0.56) by roughly half compared to the control and resulted in approximately 35% decrease in the BI (DR=0.63-0.64). All other entomological indices also showed similar and statistically significant reductions in intervention arms as compared to the control. There were no statistical differences identified between the two intervention arms, however it should be noted that the trial was not powered to detect those differences. Regardless, the lack of difference between the arms could also be due to coverage. Guppy coverage was much lower in intervention Arm 1 than in Arm 2 (~50% vs ~80%), therefore suggesting the use of PPF may have contributed to keeping entomological indicators similar to those in Arm 2.

Although none of the mosquito pools were found to be positive for dengue virus, all the positive and negative controls performed as expected. Additionally, a model used to simulate the process of mosquito sampling, pooling, and virus testing and found that mosquito infection rates commonly underestimate the prevalence of arbovirus infection in mosquito populations [45]. This suggests that in our trial either 1) the minimum infection rate found was the true rate in the population, 2) there was some degradation of RNA which resulted in untrue rates (despite proper cold chain management), or 3) the amount of virus in the pools was below the detection threshold.

It was observed that adherence to guppies was high (70-80%) and consistent when only one intervention requiring behavior change (guppies) was assigned. In the intervention arm with guppies and PPF adherence to one intervention was highest when focused health education messages were given on that intervention specifically (e.g. guppy coverage in March was highest when guppy use was emphasized and lowest in December to February when PPF usage was emphasized). Similar dynamics have been found with the use of other vector control tools. A recent review concluded that, when applied as a single intervention, temephos was found to be effective at suppressing entomological indices. However the same effect was not present when applied in combination with other interventions [46]. This suggests that unfortunately no single vector control intervention may be enough to reach elimination of dengue and using multiple interventions which require behavior change may reduce individual intervention effectiveness. Some studies have suggested combining imperfect vector control with an imperfect medium-high efficacy vaccine could be more efficacious and cost-effective way to reduce dengue cases [47,48].

The results of the KAP survey showed very high baseline knowledge levels which may have resulted from the high number of cases in the study site and from previous government-led anti-dengue efforts in these areas. The knowledge that dengue is transmitted by *Aedes* mosquitos rose to 100% of respondents by the end of the intervention, however even that was not statistically significant between baseline and endline surveys. Similarly high levels of knowledge on other dengue topics was found in the baseline survey and reported earlier [49]. Interestingly, self-reported vector control practices did not match observed practices recorded in the surveys, nor was a correlation found between knowledge and observed practices either [49]. Therefore, an education campaign regarding dengue prevention in this setting with high knowledge levels is unlikely to have any significant effect on practices unless it is incorporated in a more comprehensive strategy for behavioral change (e.g. use of the COMBI method). In addition, to bridge the knowledge-practice gap, there is a need to create an enabling environment at the household, community and health facility level to follow the required behaviors. For example, the vector control knowledge will not be enough until they have a continuous supply of the recommended interventions (e.g. guppies, PPF, *Btĩ)* in order to follow the recommended behaviors.

In previously reported Focus Group Discussions (FGDs) and In-Depth Interviews (IDIs), nearly all participants perceived that the interventions resulted in a reduction in *Aedes* mosquitos (both adults and immatures) and dengue cases [50]. Participants showed high demand for both interventions (guppies and PPF) and were willing to pay between 100-500 riel (0.03-0.13 USD). In addition, several participants began rearing guppies in their home for their personal use, for the children to play with, and to possibly sell in the market. The presence of larvae in the water despite the use of PPF was a source of concern for some participants, although this was overcome in most cases with proper health education through health volunteers. Interpersonal communication through health volunteers was the most preferred method of transmitting prevention messages. Together the entomological, KAP, and qualitative results suggest that the interventions were efficacious and accepted by the community.

However, there is always a need to balance potential benefits and harms of any intervention. Following the recent Zika outbreaks in 2015-2016, there were two groups of ecologists that noticed public health authorities utilizing non-native larvivorous fish (including guppies) in *Aedes* control [51,52]. Both of these groups wrote opinion pieces that gave three strong messages; 1) the use of larvivorous fish in vector control is not effective, 2) the chances of accidental guppy introduction into local ecosystems are large, and 3) that guppies can easily establish populations and damage these aquatic ecosystems. The first point is contradicted by studies which were available at the time, as well as by the current trial [17,18,28]. However, regarding the other points, guppies are indeed known to be highly plastic and acclimate to new environments [53]. For example, as far back as 1963 guppies have been highly effective in *Culex* control in highly polluted ground pools and waterways in Bangkok, Yangon, and Taipei [54]. In one study it was postulated that female guppies are capable of routinely establishing new populations in mesocosms, and that over 80% of these populations persist for at least two years [55]. Therefore, the key question is what is the ecological impact of guppies being accidentally released into the environment? Despite the strong statements made in the opinion pieces, the underlying evidence seems to be weaker than implied with most introductions made before proper baseline assessments were completed. Studies have shown some effects of guppies on resident fish densities in lab conditions [56,57], and nitrogen levels in water [58–60], however the extent of these effects across the ecosystem - especially in areas where introduction and naturalization took place many decades ago (such as Cambodia) - are far from settled. A book on evolutionary ecology of the guppy noted that in regards to the impact of exotic guppies “the literature is scant, and the area ripe for research” [53]. The author also noted that manner in which introduced fish species impact native assemblages is incompletely understood, and that issues such as anthropogenic changes to the habitat, such as rise in water temperature, could favor introduced over native species [53].

Measures available to control programs to mitigate the risks of introduction include; 1) restricting breeding sites to areas which can be locked and controlled by the breeders; 2) only distributing fish to key containers in at-risk areas and away from lakes and streams); 3) only distributing male fish to avoid breeding after accidental release by households; or 4) evaluating which indigenous larvivorous fish exist that have similar predation behaviors to guppies and consider their use. It should be noted that male guppies have been found to consume less larvae than males (123 per day compared to 74 per day) [17], however that consumption rate was more than enough to clear the main breeding jars in Cambodia.

In addition to concerns on accidental release of guppies to the environment, some lab experiments have raised the possibility that putting guppies in containers used for drinking water could increase *Escherichia coli* and other bacteria [61]. However, a recent study (Sidavong et al., submitted manuscript) found the addition guppy fish in Lao and Cambodia made no significant difference to high pre-existing baseline levels of contamination. Therefore, the authors concluded that any contaminating effect may be insignificant when compared with the potential for reducing dengue fever cases and advocated for the inclusion of advice on safe water use to be included in any behavior change communication programs for guppy introduction.

This study has several limitations. The most important of which is the absence of a primary outcome directly related to dengue incidence rather than an entomological one. Finding the appropriate metric to measure disease impact is bedeviled by the effect of human movement on patterns of transmission, and the pronounced temporal and spatial heterogeneity in transmission, which will necessitate very large cluster-randomized study designs [62,63]. We considered passive surveillance for dengue with rapid diagnostic tests in HCs. Although sensitivity among currently available tests was considered acceptable for routine clinical diagnostics [64] it was not considered high enough for seroconversion studies and no studies were identified that had used rapid diagnostics to estimate seroprevalence. Therefore, more expensive and labor intensive efforts were preferable, such as cohort studies or capture-recapture methods (which have their own limitations[65]) to estimate the true number of cases, or using a more sensitive diagnostic tool such as RT-PCR. However, due to budget limitations it was not possible to employ them. Additionally, unpublished data from a recent cohort study in the proposed districts suggest that, given similar number of cases during this study timeframe, and the resources available to the current project, there would not be enough statistical power to show an impact of the likely size on case numbers [31]. Therefore, the endpoint chosen was the density of adult *Aedes* mosquitoes, which are on the causal pathway to disease.

Nevertheless, determining the effect of an entomological outcome on dengue transmission is difficult. Multiple studies in Cuba have suggested that a BI of greater than five can be used to predict dengue transmission, although they note that their results can probably not be extrapolated to areas were dengue transmission is endemic [66,67]. A recent study from Peru did show a statistically significant association between 12-month longitudinal data on *Aedes aegypti* abundance (1.01-1.30) and categorical immature indices (1.21-1.75) on risk ratios dengue virus seroconversion (over six months) [66]. However, even the existence of an association remains less clear across geographies, and what the strength of that association would be in Cambodia (with much higher incidence rates) remains difficult to quantify. These efforts are frustrated by the many intersecting factors which determine dengue infection in communities including the probability of infecting and being infected by a mosquito bite, the duration of infection, treatment-seeking behavior, the risk of fever, which serotypes are present, acquired immunity in the host, coverage of interventions and background prevalence of dengue infections. The availability of quality data for each of these factors is limited in most tropical countries where the infection rates are highest.

Additional entomological limitations include only having one data collection point in each season, and no measure in the change of parity rate of adult females. The indoor resting collection of *Aedes* adult mosquitoes is subject to many challenges including: (i) individual collector performance & efficiency; (ii) density being time dependent; (iii) and housing conditions, architecture, objects, etc. Another possible source of bias is not having data collectors blind to the intervention; however, in this case it was unavoidable as data collection teams were able to see the fish in the containers which they sample. Additionally, as these data are being collected within one province in Cambodia generalizability could be a concern. However, it is likely that the result of this trial could be generalizable to areas with similar ecology and mosquito densities within the country and in neighboring countries.

In conclusion, the results from this trial indicate that the interventions resulted in a statistically significant reduction in immature and adult *Aedes* mosquito density when compared to the control. There were no statistical differences identified between intervention arms, although lower guppy coverage in intervention arm two suggests that PPF did help keep mosquito densities low. Data from the KAP and qualitative assessments showed that the interventions were accepted by communities and that they were willing to pay for them. The extremely low cost of including guppy rearing in community-based health structures along with the effectiveness demonstrated here suggests guppies should be considered as a vector control tool as long as the benefits outweigh any potential environmental concerns. PPF was also highly accepted and preferred over current vector control tools used in Cambodia, however product costs and availability are still unknown. The qualitative assessment suggests that a context specific and well-informed COMBI and community engagement by giving an active role to communities is the key to the successful dengue control. Additional studies could be done to confirm these results and explore the effect of the interventions is different ecological conditions.

## Supporting information captions

**S1 Figure:**
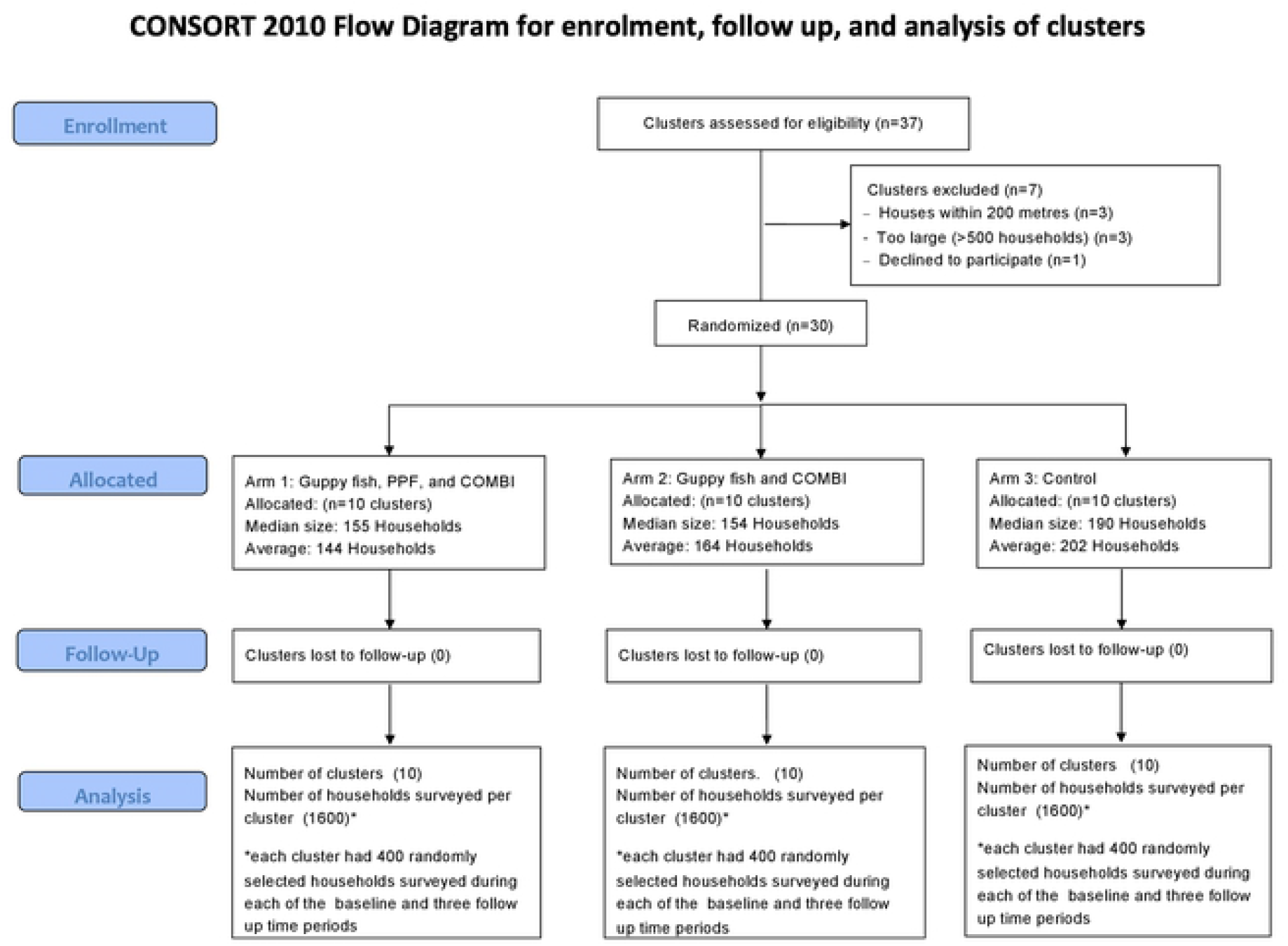
CONSORT 2010 Flow Diagram

S1 Table: CONSORT 2010 checklist of information to include when reporting a randomized trial

S2 Table: Remaining Secondary Outcome Tables

S1 Dataset: Entomology data at cluster level used for analysis

## References

1. Seng CM, Setha T, Nealon J, Socheat D, Nathan MB. Six months of *Aedes* aegypti control with a novel controlled-release formulation of pyriproxyfen in domestic water storage containers in Cambodia. Southeast Asian J Trop Med Public Heal. 2008;39: 822–826. Available: http://www.ncbi.nlm.nih.gov/pubmed/19058575

2. Vannice KS, Roehrig JT, Hombach J. Next generation dengue vaccines: A review of the preclinical development pipeline. Vaccine. 2015; doi:10.1016/j.vaccine.2015.09.053

3. Alphey L, Mckemey A, Nimmo D, Oviedo MN, Lacroix R, Matzen K, et al. Genetic control of *Aedes* mosquitoes. Pathog Glob Health. 2013; doi:10.1179/2047773213Y.0000000095

4. Franz AWE, Clem RJ, Passarelli AL. Novel genetic and molecular tools for the investigation and control of dengue virus transmission by mosquitoes. Current Tropical Medicine Reports. 2014. doi:10.1007/s40475-013-0007-2

5. Ye YH, Carrasco AM, Frentiu FD, Chenoweth SF, Beebe NW, van den Hurk AF, et al. Wolbachia reduces the transmission potential of dengue-infected *Aedes aegypti*. PLoS Negl Trop Dis. 2015; doi:10.1371/journal.pntd.0003894

6. Turelli M, Barton NH. Deploying dengue-suppressing Wolbachia: Robust models predict slow but effective spatial spread in *Aedes aegypti*. Theor Popul Biol. 2017; doi:10.1016/j.tpb.2017.03.003

7. Bowman LR, Donegan S, McCall PJ. Is dengue vector control deficient in effectiveness or evidence?: systematic review and meta-analysis. PLoS Negl Trop Dis. 2016; doi:10.1371/journal.pntd.0004551

8. Huy R, Buchy P, Conan A, Ngan C, Ong S, Ali R, et al. National dengue surveillance in Cambodia 1980-2008: epidemiological and virological trends and the impact of vector control. Bull World Heal Organ. 2010;88: 650–657. doi:10.2471/BLT.09.073908

9. Vong S, Goyet S, Ly S, Ngan C, Huy R, Duong V, et al. Under-recognition and reporting of dengue in Cambodia: A capture-recapture analysis of the National Dengue Surveillance System. Epidemiol Infect. 2012; doi:10.1017/S0950268811001191

10. Wichmann O, Yoon IK, Vong S, Limkittikul K, Gibbons R V., Mammen MP, et al. Dengue in Thailand and Cambodia: An assessment of the degree of underrecognized disease burden based on reported cases. PLoS Negl Trop Dis. 2011; doi:10.1371/journal.pntd.0000996

11. Seng CM, Setha T, Nealon J, Chantha N, Socheat D, Nathan MB. The effect of long-lasting insecticidal water container covers on field populations of *Aedes aegypti* (L.) mosquitoes in Cambodia. J Vector Ecol. 2008;33: 333–341. Available: http://www.ncbi.nlm.nih.gov/pubmed/19263854

12. Khun S, Manderson LH. Abate distribution and dengue control in rural Cambodia. Acta Trop. 2007;101: 139–146. doi:10.1016/j.actatropica.2007.01.002

13. Boyer S, Lopes S, Prasetyo D, Hustedt J, Sarady AS, Doum D, et al. Resistance of *Aedes aegypti* (Diptera: Culicidae) populations to Deltamethrin, Permethrin, and Temephos in Cambodia. Asia-Pacific J Public Heal. 2018; doi:10.1177/1010539517753876

14. Polson KA, Curtis C, Seng CM, Olson JG, Chantha N, Rawlins SC. Susceptibility of two cambodian population of *Aedes aegypti* mosquito larvae to temephos during 2001. Dengue Bull. 2001;25: 79–83.

15. Seng CM, Setha T, Chanta N, Socheat D, Guillet P, Nathan MB. Inhibition of adult emergence of *Aedes aegypti* in simulated domestic water-storage containers by using a controlled-release formulation of pyriproxyfen. J Am Mosq Control Assoc. United States; 2006;22: 152–154. doi:10.2987/8756-971X(2006)22[152:IOAEOA]2.0.CO;2

16. Setha T, Chantha N, Socheat D. Efficacy of *Bacillus thuringiensis israelensis*, VectoBac WG and DT, formulations against dengue mosquito vectors in cement potable water jars in Cambodia. Southeast Asian J Trop Med Public Heal. 2007/06/02. 2007;38: 261–268.

17. Seng CM, Setha T, Nealon J, Socheat D, Chantha N, Nathan MB, et al. Community-based use of the larvivorous fish Poecilia reticulata to control the dengue vector *Aedes aegypti* in domestic water storage containers in rural Cambodia. J Vector Ecol. 2008/08/14. 2008;33: 139–144. doi:10.3376/1081-1710(2008)33[139:CUOTLF]2.0.CO;2

18. World Health Organization. Managing regional public goods for health: community-based dengue vector control [Internet]. Mandaluyong City, Philippines; 2013. Available: https://www.adb.org/sites/default/files/publication/30167/community-based-dengue-vector-control.pdf

19. Woosome K. Tiny predator has big role curbing dengue. The Cambodia Daily. 2004. Available: https://www.cambodiadaily.com/archives/tiny-predator-has-big-role-curbing-dengue-37552/

20. World Health Organization. Report of the fourth WHOPES working group meeting. Review of IR3535, KBR3023, (RS)-methoprene 20% EC and pyriproxyfen 0.5% GR. [Internet]. Geneva: World Health Organization; 2000. Available: https://apps.who.int/iris/handle/10665/66683

21. Sai-Zaw-Min-Oo, Sein-Thaung, Yan-Naung-Maung-Maung, Khin-Myo-Aye, Zar-Zar-Aung, Hlaing-Myat-Thu, et al. Effectiveness of a novel long-lasting pyriproxyfen larvicide (SumiLarv^®^2MR) against *Aedes* mosquitoes in schools in Yangon, Myanmar. Parasites and Vectors. 2018; doi:10.1186/s13071-017-2603-9

22. Suaya JA, Shepard DS, Chang MS, Caram M, Hoyer S, Socheat D, et al. Cost-effectiveness of annual targeted larviciding campaigns in Cambodia against the dengue vector *Aedes aegypti*. Trop Med Int Heal. 2007/09/19. 2007;12: 1026–1036. doi:10.1111/j.1365-3156.2007.01889.x

23. Azmawati MN, Aniza I, Ali M. Evaluation of Communication for Behavioral Impact (COMBI) program in dengue prevention: a qualitative and quantitative study in Selangor, Malaysia. Iran J Public Heal. 2013;42: 538–539. Available: http://www.ncbi.nlm.nih.gov/pubmed/23802114

24. Al-Muhandis N, Hunter PR. The value of educational messages embedded in a community-based approach to combat dengue fever: a systematic review and meta regression analysis. PLoS Negl Trop Dis. 2011/09/03. 2011;5: e1278. doi:10.1371/journal.pntd.0001278

25. Heintze C, Velasco Garrido M, Kroeger A. What do community-based dengue control programmes achieve? A systematic review of published evaluations. Trans R Soc Trop Med Hyg. 2006/11/07. 2007;101: 317–325. doi:10.1016/j.trstmh.2006.08.007

26. Vanlerberghe V, Toledo ME, Rodriguez M, Gomez D, Baly A, Benitez JR, et al. Community involvement in dengue vector control: cluster randomised trial. MEDICC Rev. 2010/04/15. 2010;12: 41–47.

27. Castro M, Sanchez L, Perez D, Carbonell N, Lefevre P, Vanlerberghe V, et al. A community empowerment strategy embedded in a routine dengue vector control programme: a cluster randomised controlled trial. Trans R Soc Trop Med Hyg. 2012/04/03. 2012;106: 315–321. doi:10.1016/j.trstmh.2012.01.013

28. Han WW, Lazaro A, McCall PJ, George L, Runge-Ranzinger S, Toledo J, et al. Efficacy and community effectiveness of larvivorous fish for dengue vector control. Trop Med Int Heal. 2015;20: 1239–1256. doi:10.1111/tmi.12538

29. Campbell MK, Piaggio G, Elbourne DR, Altman DG. Consort 2010 statement: Extension to cluster randomised trials. BMJ. 2012; doi:10.1136/bmj.e5661

30. Harrington LC, Scott TW, Lerdthusnee K, Coleman RC, Costero A, Clark GG, et al. Dispersal of the dengue vector *Aedes aegypti* within and between rural communities. Am J Trop Med Hyg. 2005;72: 209–220. Available: http://www.ncbi.nlm.nih.gov/pubmed/15741559

31. Hustedt J, Doum D, Keo V, Ly S, Sam BL, Chan V, et al. Determining the efficacy of guppies and pyriproxyfen (Sumilarv^®^ 2MR) combined with community engagement on dengue vectors in Cambodia: Study protocol for a randomized controlled trial. Trials. 2017; doi:10.1186/s13063-017-2105-2

32. World Health Organization. WHO Specifications and Evaluations for Public Health Pesticides - Pyriproxyfen [Internet]. 2017. Available: https://www.who.int/whopes/quality/Pyriproxyfen_eval_specs_WHO_July_2017.pdf

33. World Health Organization. Report of the Twentieth WHOPES Working Group Meeting [Internet]. 2017. Available: https://apps.who.int/iris/bitstream/handle/10665/258921/WHO-HTM-NTD-WHOPES-2017.04-eng.pdf

34. Hemming K MJ. A menu-driven facility for sample-size calculations in cluster randomized controlled trials. Stata J. 2013;13: 114–135.

35. Campbell MK, Fayers PM, Grimshaw JM. Determinants of the intracluster correlation coefficient in cluster randomized trials: the case of implementation research. Clin Trials. 2005;2: 99–107. Available: http://www.ncbi.nlm.nih.gov/pubmed/16279131

36. World Health Organzation. Dengue: guidelines for diagnosis, treatment, prevention, and control. Spec Program Res Train Trop Dis. Geneva: World Health Organization; 2009; 147. doi:WHO/HTM/NTD/DEN/2009.1

37. Tun-Lin W, Kay BH, Barnes A. The Premise Condition Index: a tool for streamlining surveys of *Aedes aegypti*. Am J Trop Med Hyg. 1995/12/01. 1995;53: 591–594. Available: https://www.ncbi.nlm.nih.gov/pubmed/8561259

38. Hustedt J, Doum D, Keo V, Ly S, Sam B, Chan V, et al. Ability of the Premise Condition Index to Identify Premises with Adult and Immature *Aedes* Mosquitoes in Kampong Cham, Cambodia. Am J Trop Med Hyg. 2020; doi:10.4269/ajtmh.19-0453

39. Sule WF, Oluwayelu DO. Analysis of culex and aedes mosquitoes in southwestern Nigeria revealed no west nile virus activity. Pan Afr Med J. 2016; doi:10.11604/pamj.2016.23.116.7249

40. Cevallos V, Ponce P, Waggoner JJ, Pinsky BA, Coloma J, Quiroga C, et al. Zika and Chikungunya virus detection in naturally infected *Aedes aegypti* in Ecuador. Acta Trop. 2018; doi:10.1016/j.actatropica.2017.09.029

41. Pierre V, Drouet MT, Deubel V. Identification of mosquito-borne flavivirus sequences using universal primers and reverse transcription/polymerase chain reaction. Res Virol. 1994; doi:10.1016/S0923-2516(07)80011-2

42. Lanciotti RS, Calisher CH, Gubler DJ, Chang GJ, Vorndam A V. Rapid detection and typing of dengue viruses from clinical samples by using reverse transcriptase-polymerase chain reaction. J Clin Microbiol. 1992;

43. Grant AM, Altman DG, Babiker AB, Campbell MK, Clemens FJ, Darbyshire JH, et al. Issues in data monitoring and interim analysis of trials. Heal Technol Assess. 2005;9: 1–238, iii-iv. Available: http://www.ncbi.nlm.nih.gov/pubmed/15763038

44. American Anthropological Association. Principles of Professional Responsibility [Internet]. 2012. Available: http://ethics.aaanet.org/category/statement/

45. Bustamante DM, Lord CC. Sources of error in the estimation of mosquito infection rates used to assess risk of arbovirus transmission. Am J Trop Med Hyg. 2010; doi:10.4269/ajtmh.2010.09-0323

46. George L, Lenhart A, Toledo J, Lazaro A, Han WW, Velayudhan R, et al. Community-effectiveness of temephos for dengue vector control: a systematic literature review. PLoS Negl Trop Dis. 2015;9: e0004006. doi:10.1371/journal.pntd.0004006

47. Fitzpatrick C, Haines A, Bangert M, Farlow A, Hemingway J, Velayudhan R. An economic evaluation of vector control in the age of a dengue vaccine. PLoS Negl Trop Dis. 2017; doi:10.1371/journal.pntd.0005785

48. Christofferson RC, Mores CN. A role for vector control in dengue vaccine programs. Vaccine. 2015; doi:10.1016/j.vaccine.2015.09.114

49. Kumaran E, Doum D, Keo V, Sokha L, Sam BL, Chan V, et al. Dengue knowledge, attitudes and practices and their impact on community-based vector control in rural Cambodia. PLoS Negl Trop Dis. 2018; doi:10.1371/journal.pntd.0006268

50. Shafique M, Lopes S, Doum D, Keo V, Sokha L, Sam B, et al. Implementation of guppy fish (Poecilia reticulata), and a novel larvicide (Pyriproxyfen) product (Sumilarv 2MR) for dengue control in Cambodia: A qualitative study of acceptability, sustainability and community engagement. PLoS Negl Trop Dis. 2019;13: e0007907. doi:10.1371/journal.pntd.0007907

51. El-Sabaawi RW, Frauendorf TC, Marques PS, Mackenzie RA, Manna LR, Mazzoni R, et al. Biodiversity and ecosystem risks arising from using guppies to control mosquitoes. Biology letters. 2016. doi:10.1098/rsbl.2016.0590

52. Azevedo-Santos VM, Vitule JRS, Pelicice FM, García-Berthou E, Simberloff D. Nonnative fish to control *Aedes* mosquitoes: A controversial, harmful tool. BioScience. 2017. doi:10.1093/biosci/biw156

53. Magurran AE. Evolutionary ecology: the Trinidadian guppy. Oxford Series in Ecology and Evolution. 2005. doi:10.1093/acprof

54. Bay EC, Self LS. Observations of the guppy, Poecilia reticulata Peters, in Culex pipiens fatigans breeding sites in Bangkok, Rangoon, and Taipei. Bull World Health Organ. 1972;

55. Deacon AE, Ramnarine IW, Magurran AE. How reproductive ecology contributes to the spread of a globally invasive fish. PLoS One. 2011; doi:10.1371/journal.pone.0024416

56. Walsh MR, Reznick DN. Influence of the indirect effects of guppies on life-history evolution in rivulus hartii. Evolution (N Y). 2010; doi:10.1111/j.1558-5646.2009.00922.x

57. Walsh MR, Reznick DN. Experimentally induced life-history evolution in a killifish in response to the introduction of guppies. Evolution (N Y). 2011; doi:10.1111/j.1558-5646.2010.01188.x

58. Holitzki TM, MacKenzie RA, Wiegner TN, McDermid KJ. Differences in ecological structure, function, and native species abundance between native and invaded Hawaiian streams. Ecol Appl. 2013; doi:10.1890/12-0529.1

59. El-Sabaawi RW, Marshall MC, Bassar RD, López-Sepulcre A, Palkovacs EP, Dalton C. Assessing the effects of guppy life history evolution on nutrient recycling: From experiments to the field. Freshw Biol. 2015; doi:10.1111/fwb.12507

60. Collins SM, Thomas SA, Heatherly T, MacNeill KL, Leduc AOHC, López-Sepulcre A, et al. Fish introductions and light modulate food web fluxes in tropical streams: A whole-ecosystem experimental approach. Ecology. 2016; doi:10.1002/ecy.1530

61. Chadee DD. Bacterial pathogens isolated from guppies (poecilia reticulata) used to control *Aedes aegypti* in trinidad. Trans R Soc Trop Med Hyg. 1992; doi:10.1016/0035-9203(92)90194-H

62. Van Breukelen GJP, Candel MJJM. Calculating sample sizes for cluster randomized trials: We can keep it simple and efficient! J Clin Epidemiol. 2012; doi:10.1016/j.jclinepi.2012.06.002

63. Hayes RJ, Moulton LH. Cluster Randomized Trials. Florida: Chapman & Hall; 2009.

64. Hunsperger EA, Yoksan S, Buchy P, Nguyen VC, Sekaran SD, Enria DA, et al. Evaluation of commercially available diagnostic tests for the detection of dengue virus NS1 antigen and anti-dengue virus IgM antibody. PLoS Negl Trop Dis. 2014;8: e3171. doi:10.1371/journal.pntd.0003171

65. Tilling K. Capture-recapture methods--useful or misleading? Int J Epidemiol. 2001;30: 12–14. Available: http://www.ncbi.nlm.nih.gov/pubmed/11171841

66. Sanchez L, Vanlerberghe V, Alfonso L, Marquetti MDC, Guzman MG, Bisset J, et al. *Aedes aegypti* larval indices and risk for dengue epidemics. Emerg Infect Dis. 2006; doi:10.3201/eid1205.050866

67. Sanchez L, Cortinas J, Pelaez O, Gutierrez H, Concepción D, Van Der Stuyft P. Breteau Index threshold levels indicating risk for dengue transmission in areas with low *Aedes* infestation. Trop Med Int Heal. 2010; doi:10.1111/j.1365-3156.2009.02437.x

